## Supplemental material for "Dynamic effects of genetic variation on gene expression revealed following hypoxic stress in cardiomyocytes"

### Table of contents

### Figures

Fig.S1: Purity of cardiomyocyte cultures.

Fig.S2: iPSC-CMs elicit a response to cellular stress.

Fig.S3: Numbers of RNA-seq reads are similar across conditions.

Fig.S4: Correlation of read counts across samples.

Fig.S5: Correlation of gene expression measurements across samples.

Fig.S6: Cardiomyocyte marker genes are expressed in iPSC-CMs.

Fig.S7: RNA-seq samples cluster by oxygen level and individual.

Fig.S8: Hypoxia and re-oxygenation induces a gene expression response.

Fig.S9: eQTL and dynamic eQTL identification.

Fig.S10: Overlap of response genes and eGenes with tissue eQTLs.

Fig.S11: Overlap of response genes and eGenes with TFs.

Fig.S12: Numbers of ATAC-seq reads are similar across conditions.

Fig.S13: ATAC-seq library quality control.

Fig.S14: ATAC-seq libraries cluster by individual and treatment.

Fig.S15: Identification of differentially accessible chromatin regions.

Fig.S16: Identification of caQTLs.

Fig.S17: DNA methylation array quality control.

Fig.S18: DNA methylation levels cluster by individual.

Fig.S19: DNA methylation levels are stable across conditions.

Fig.S20: Power to detect eQTLs and false positive rate to call dynamic eQTLs.

**Tables**

Table S1: Numbers of mapped RNA-seq and ATAC-seq reads.

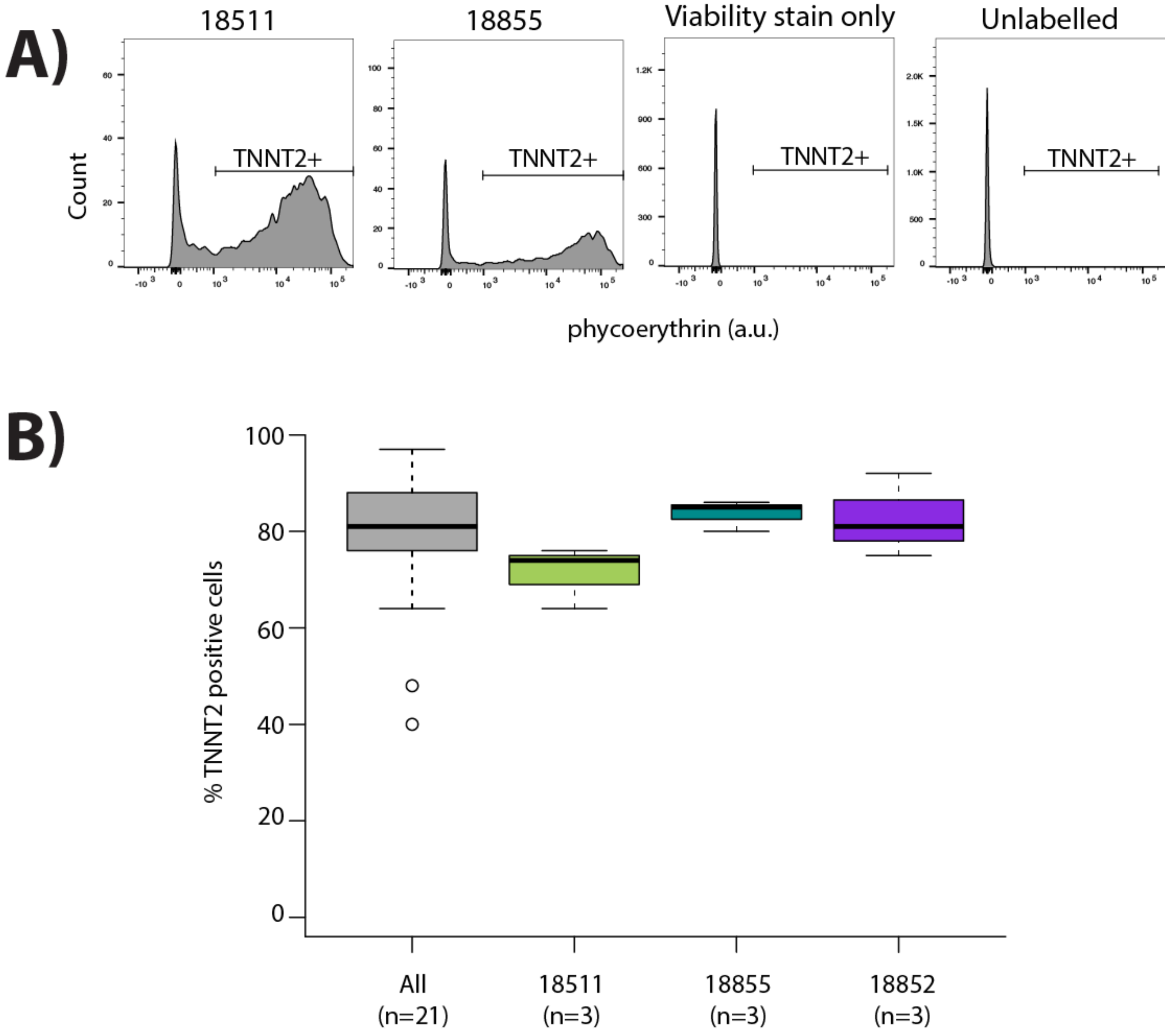

**Fig.S1: Purity of cardiomyocyte cultures. A)** Representative images from flow cytometry experiments for iPSC-derived cardiomyocytes that are labeled with cardiac troponin T (TNNT2) antibody and viability stain, and negative controls cells that are only labeled with viability stain, or are not labeled with either viability stain or TNNT2 antibody. **B)** Proportion of cells that are positive for expression of TNNT2 across all samples (All) and segregated by individuals which have replicate samples (18511, 18855, 18852).

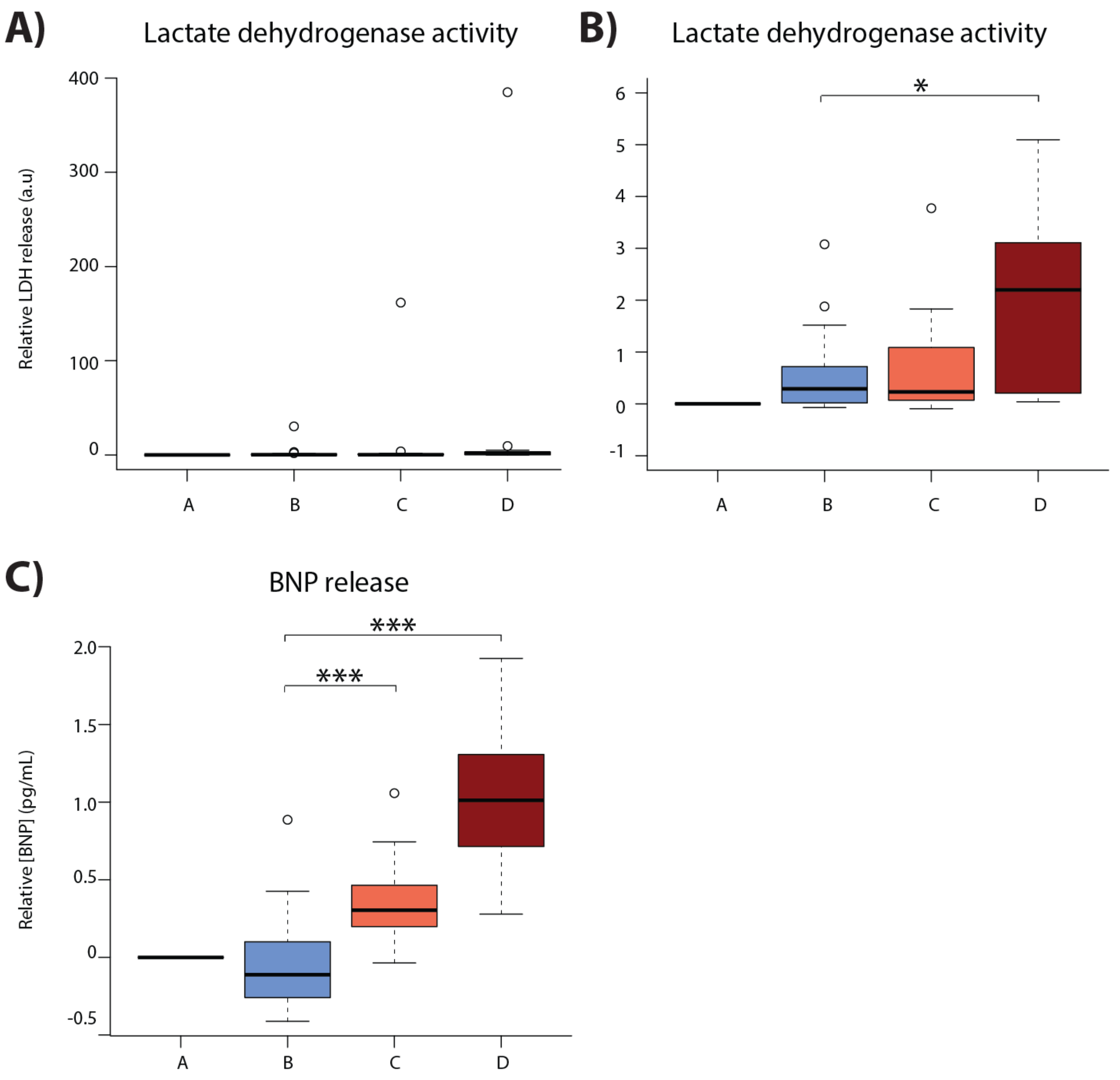

**Fig.S2: iPSC-CMs elicit a response to cellular stress. A)** Lactate dehydrogenase activity assays performed on cell culture media. **B)** Lactate dehydrogenase activity assays excluding the outlier individual. **C)** BNP ELISA assays performed on cell culture media. One representative individual for individuals with replicate data is plotted. Asterisk denotes a significant difference between conditions ($P^* < 0.05$, $P^{***} < 0.0005$).

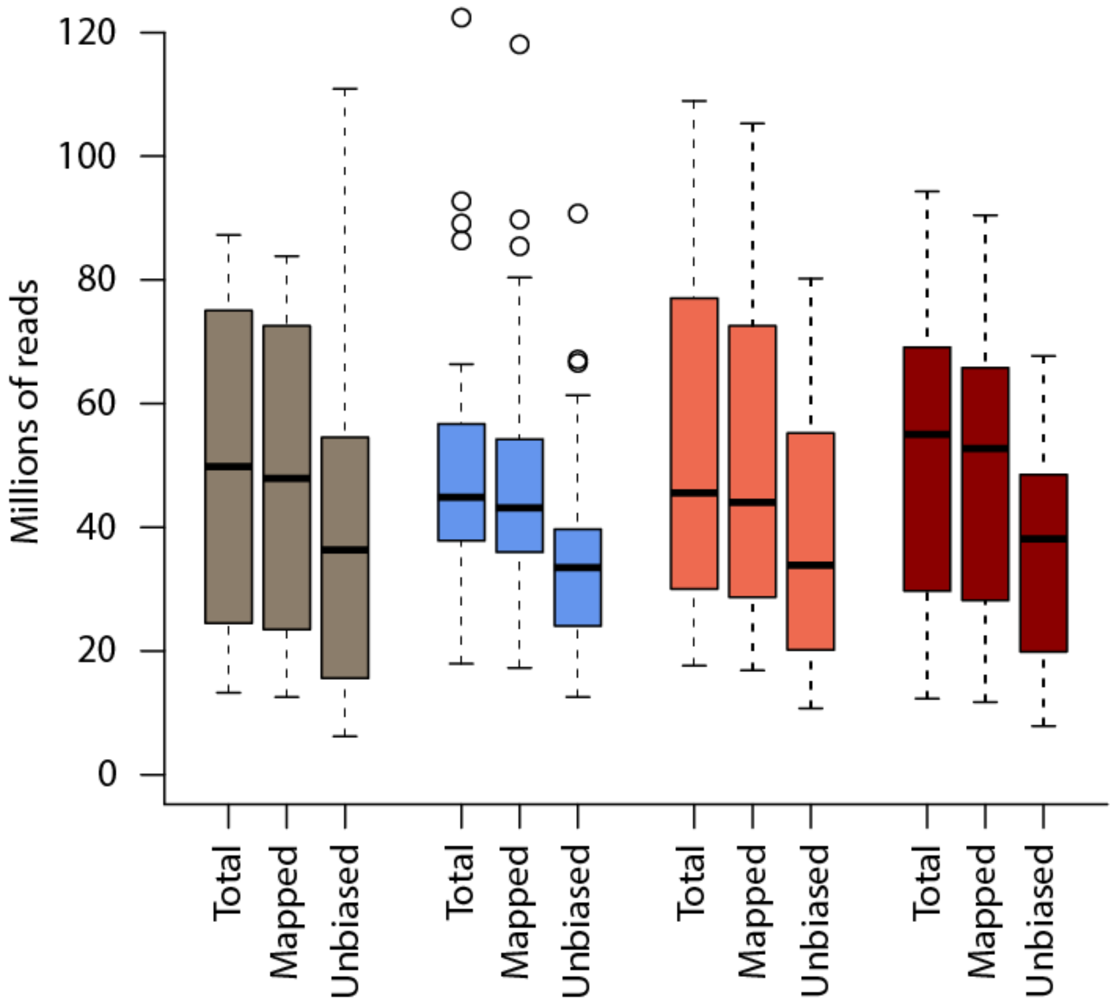

**Fig.S3: Numbers of RNA-seq reads are similar across conditions.** The total number of reads (Total), the number of reads that map to the human genome (Mapped), and the number of reads that pass the WASP re-mapping step (Unbiased) are shown for each sample within each of the four conditions (A: brown, B: blue, C: coral, D: red).

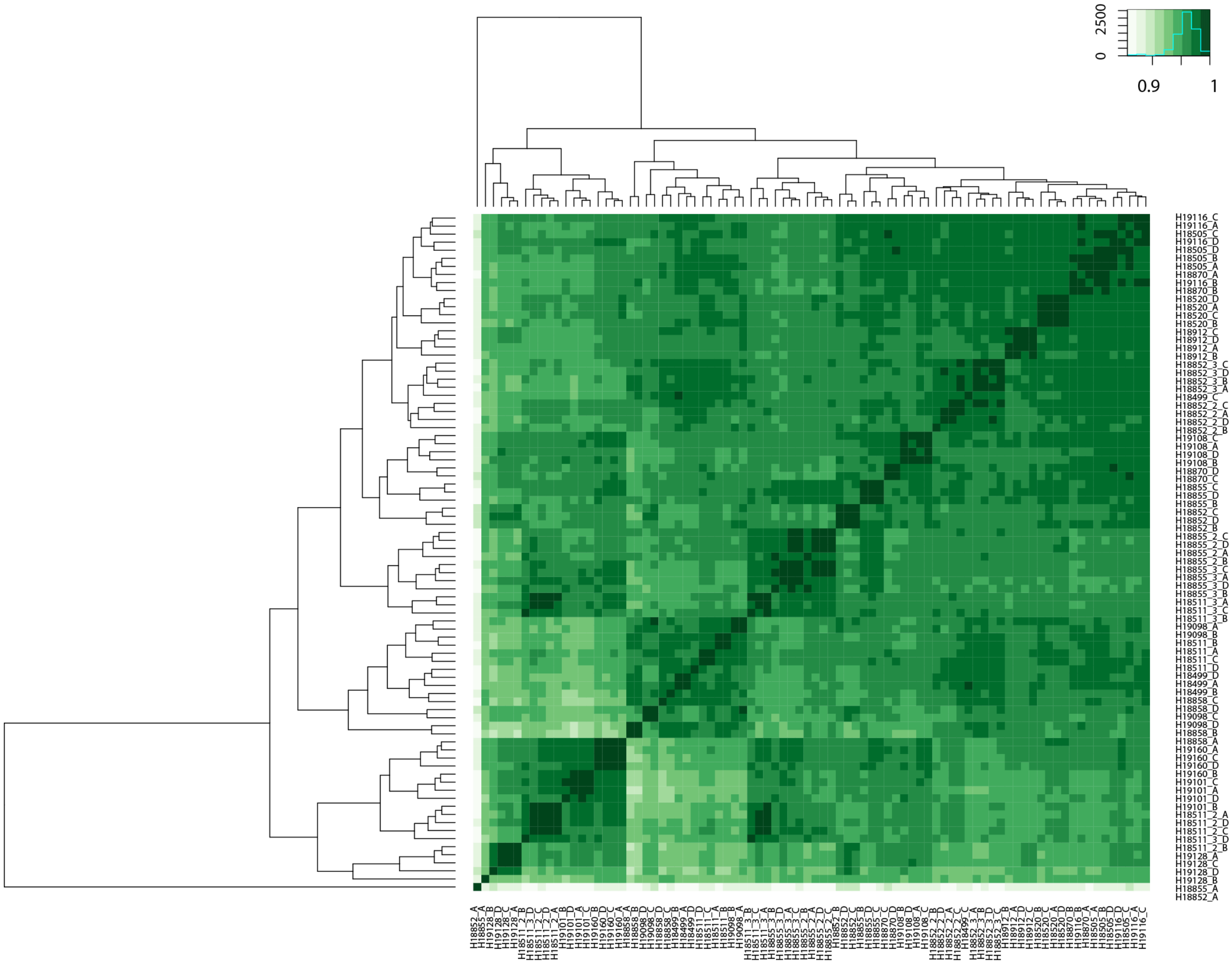

**Fig.S4: Correlation of read counts across samples.** Spearman correlation of RNA-seq read counts between all pairs of samples.

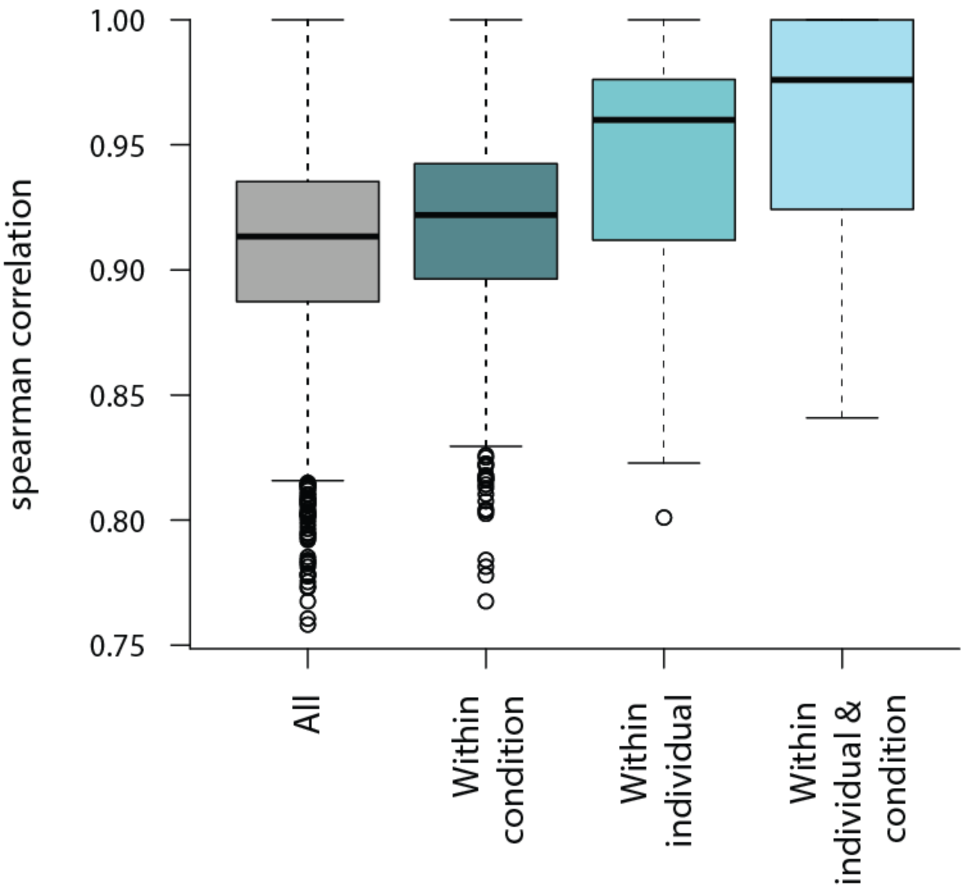

**Fig.S5: Correlation of gene expression measurements across samples.** Spearman correlation between all samples, samples within the same condition, samples from the same individual, and samples from the same condition and individual.

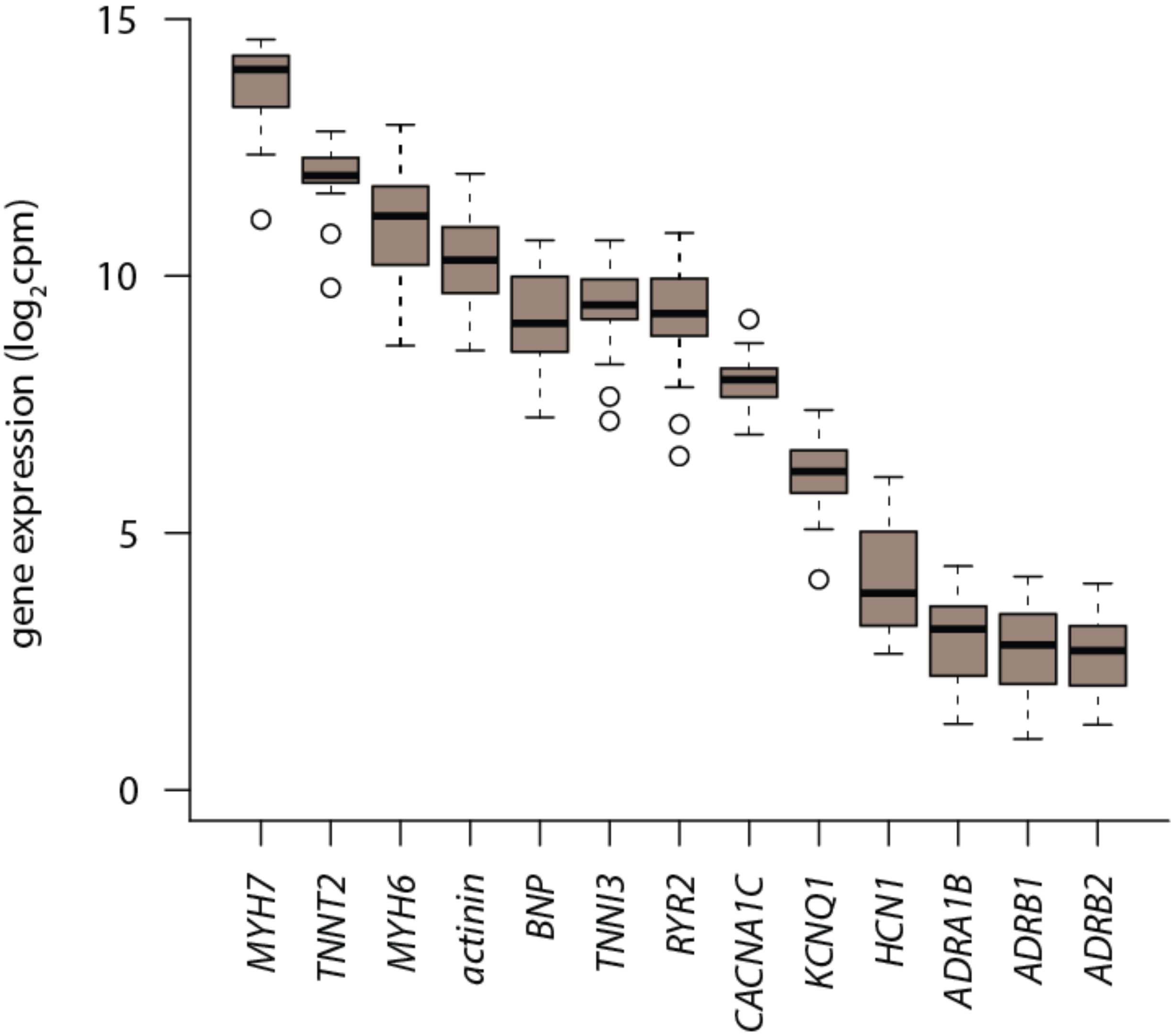

**Fig.S6: Cardiomyocyte marker genes are expressed in iPSC-CMs.** $Log_2$cpm expression levels for a panel of genes known to be expressed in cardiomyocytes. Values in Condition A (normoxia) are plotted.

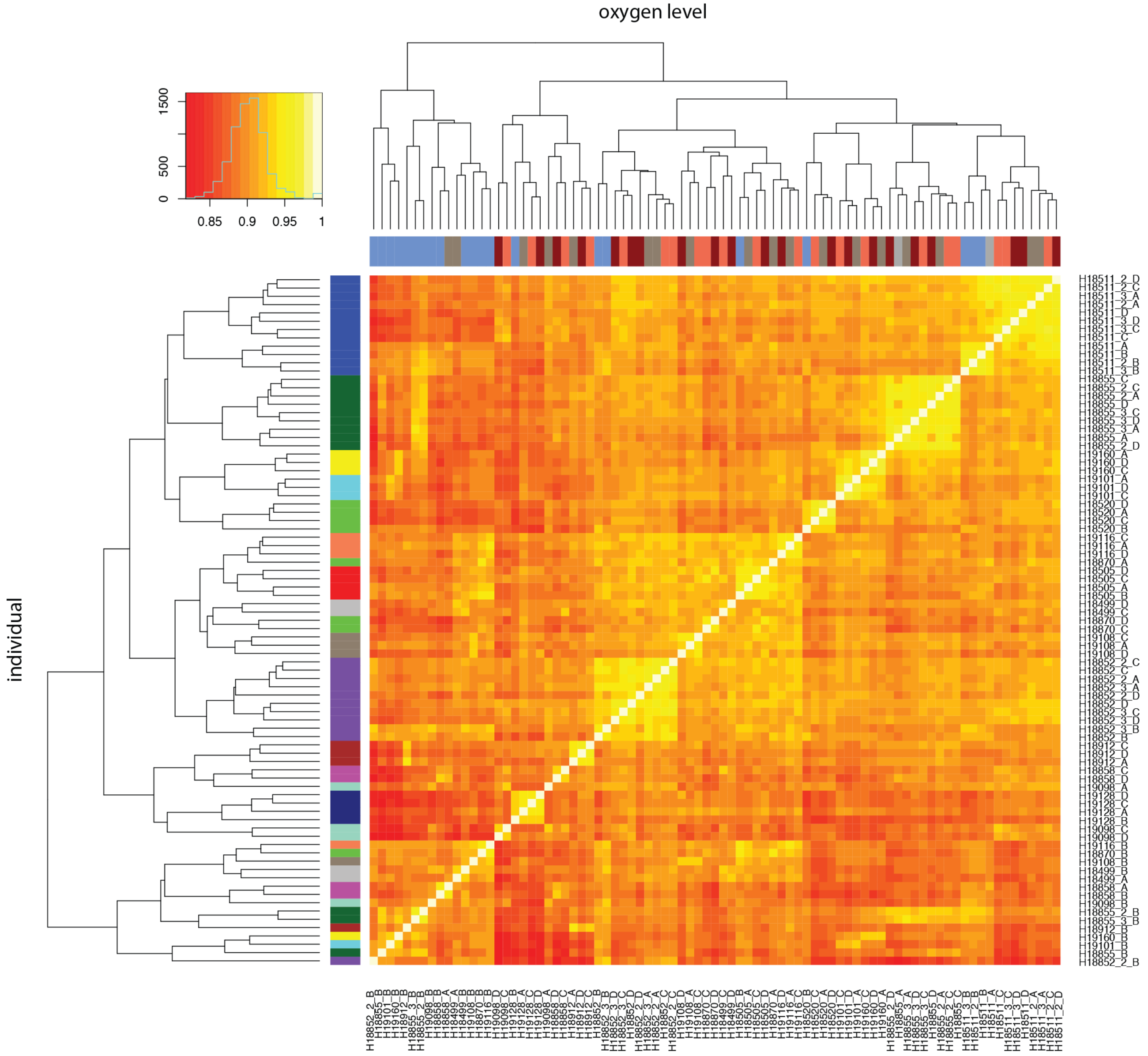

**Fig.S7: RNA-seq samples cluster by oxygen level and individual.** Spearman correlation of RUVs-normalized $\log_2$cpm expression levels between all pairs of samples. X-axis: oxygen level: normoxia at 10% oxygen (condition A: brown), 1% oxygen for 6 hours (condition B: blue), re-oxygenation to 10% oxygen for 6 hours (condition C: coral), re-oxygenation to 10% oxygen for 24 hours (condition D: red). Y-axis: individual. All samples from the same individual are represented by the same color.

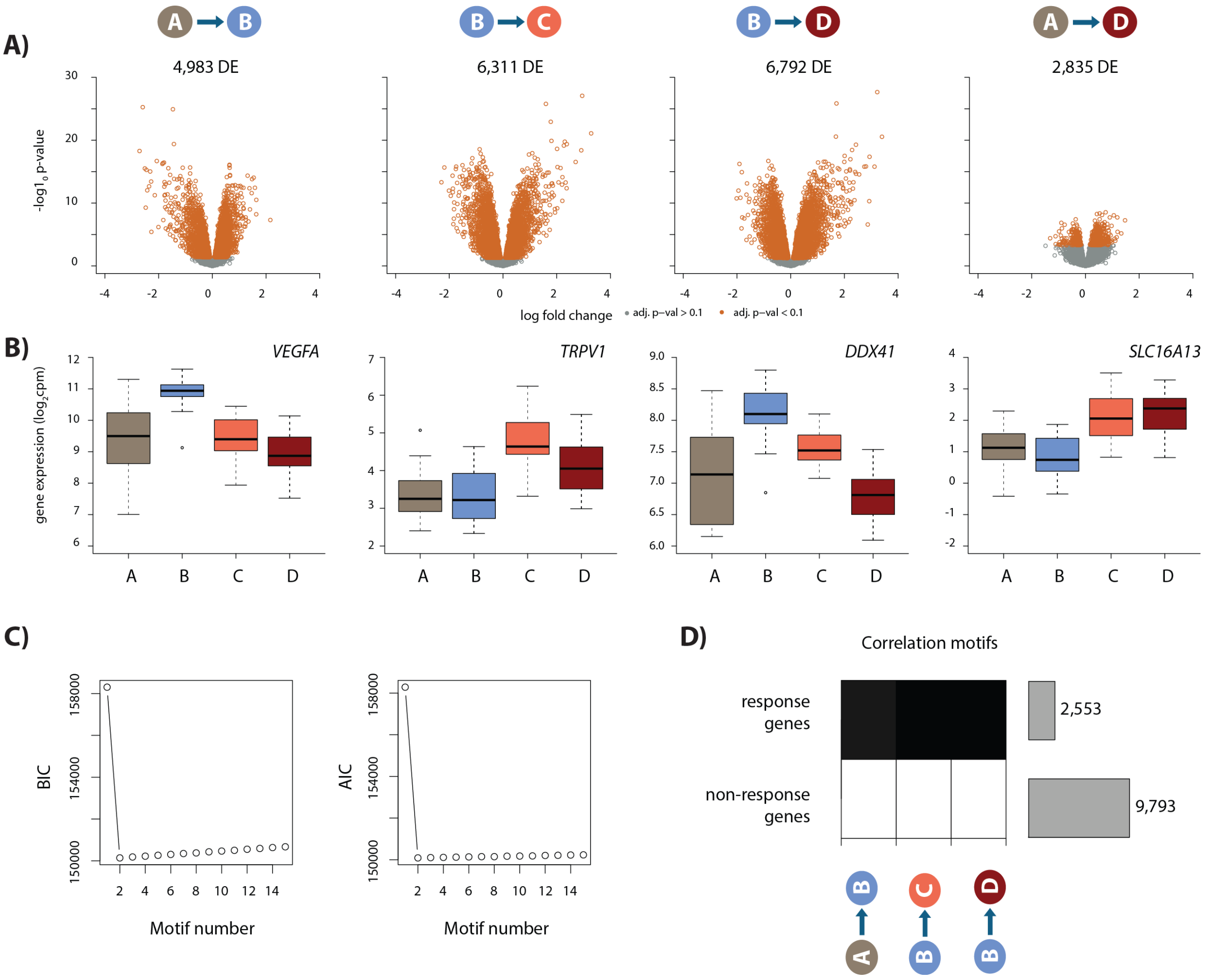

**Fig.S8: Hypoxia and re-oxygenation induces a gene expression response. A)** Volcano plots representing genes that are differentially expressed (DE) between pairs of conditions (orange dots). **B)** Examples of genes that are differentially expressed between each pair of conditions. **C)** Bayesian information criterion (BIC) and Akaike information criterion (AIC) at increasing numbers of Cormotif correlation motifs. **D)** The two correlation motifs identified by Cormotif. Color represents the posterior probability of genes being differentially expressed between pairs of conditions. Genes with a $p > 0.5$ across all tests are defined as "response genes" and those with $p < 0.5$ as "non-response genes". The number of genes associated with each motif is plotted on the right.

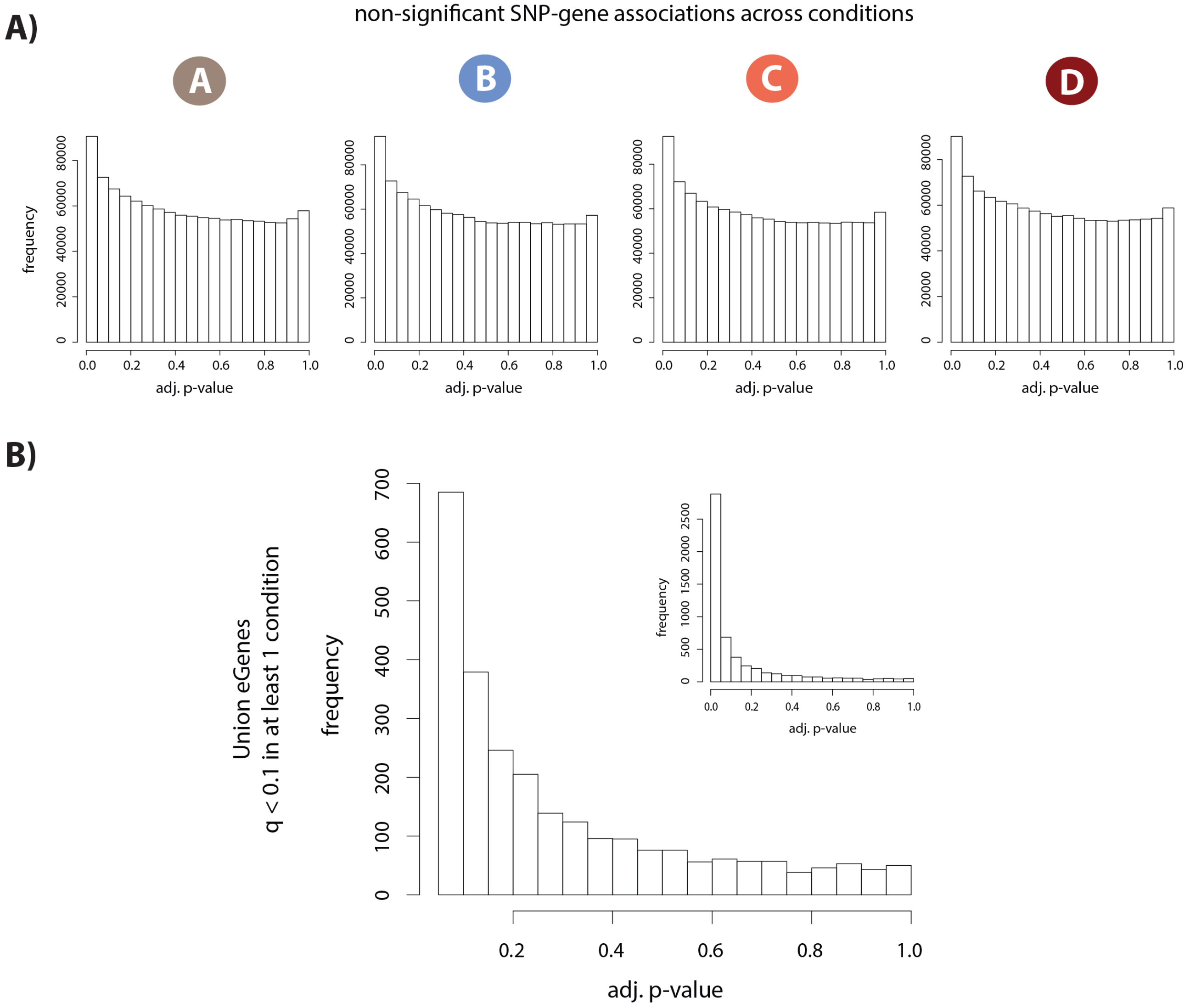

**Fig.S9: eQTL and dynamic eQTL identification. A)** Distribution of permutation-adjusted p-values in each condition for all tested SNP-gene pairs that are not significantly associated (q > 0.1). **B)** Distribution of permutation-adjusted p-values for all non-significant SNP-gene associations (q < 0.1 in at least one condition, and q > 0.1 in at least one condition) where all p-values greater than 0.05 are plotted. Inset: distribution of all p-values including those < 0.05.

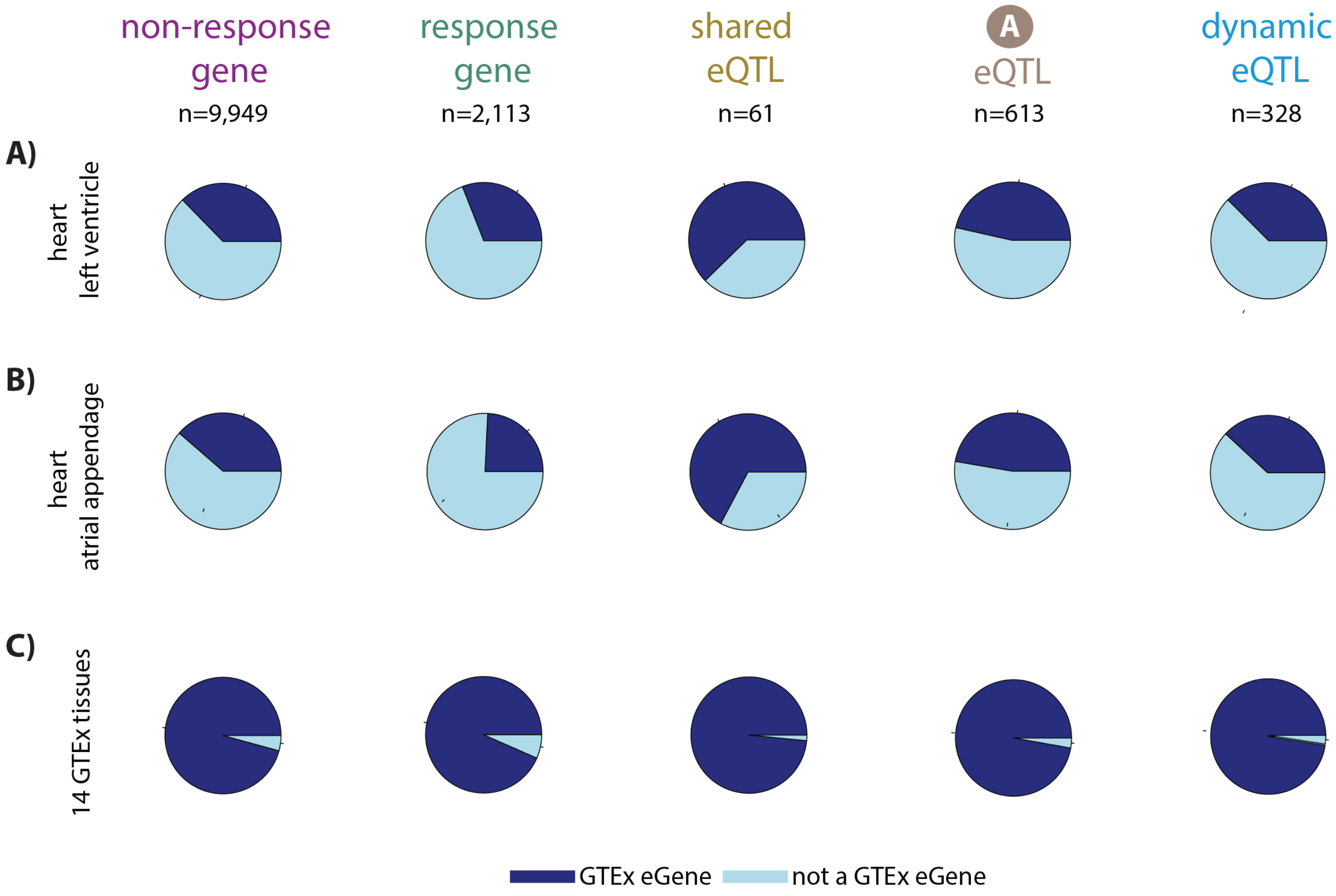

**Fig.S10: Overlap of response genes and eGenes with tissue eQTLs. A)** Enrichment of eQTLs from GTEx heart left ventricle tissue (GTEx consortium, 2017) in non-response genes, response genes, shared eQTLs, baseline eQTLs, and dynamic eQTLs. **B)** Enrichment of eQTLs from GTEx heart atrial appendage in the five gene categories. **C)** Enrichment of eQTLs from 14 GTEx tissues in our gene categories.

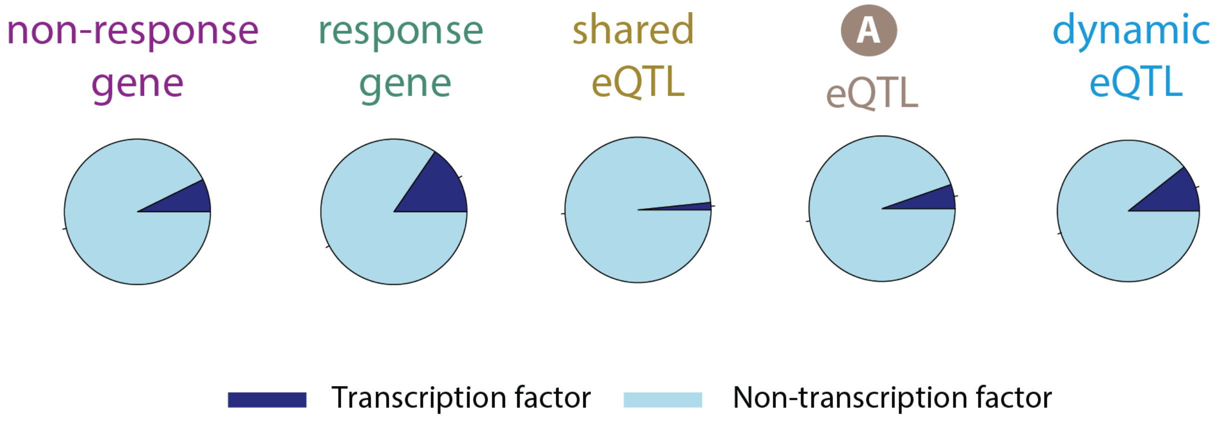

**Fig.S11: Overlap of response genes and eGenes with TFs.** Enrichment of curated transcription factors (Lambert *et al*., 2018) in our non-response genes, response genes, and eGenes.

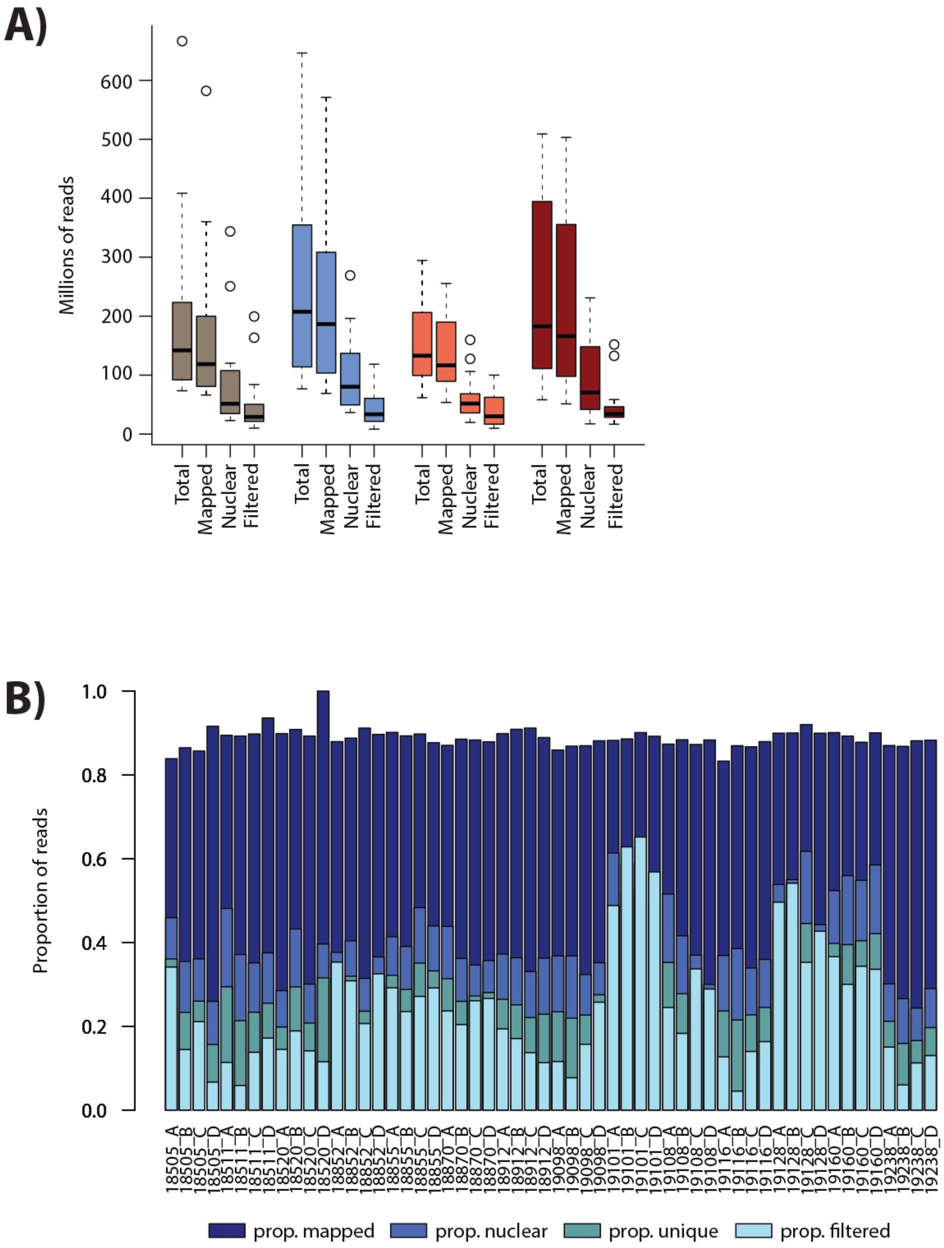

**Fig.S12: Numbers of ATAC-seq reads are similar across conditions. A)** The total number of reads (Total), the number of reads that map to the human genome (Mapped), the number of reads that map to the nuclear genome (Nuclear), and the number of reads that pass the WASP re-mapping and quality filtering step (Filtered) are shown for each sample within each of the four conditions (A: brown, B: blue, C: coral, D: red). **B)** The proportion of reads in each mapping category for each sample.

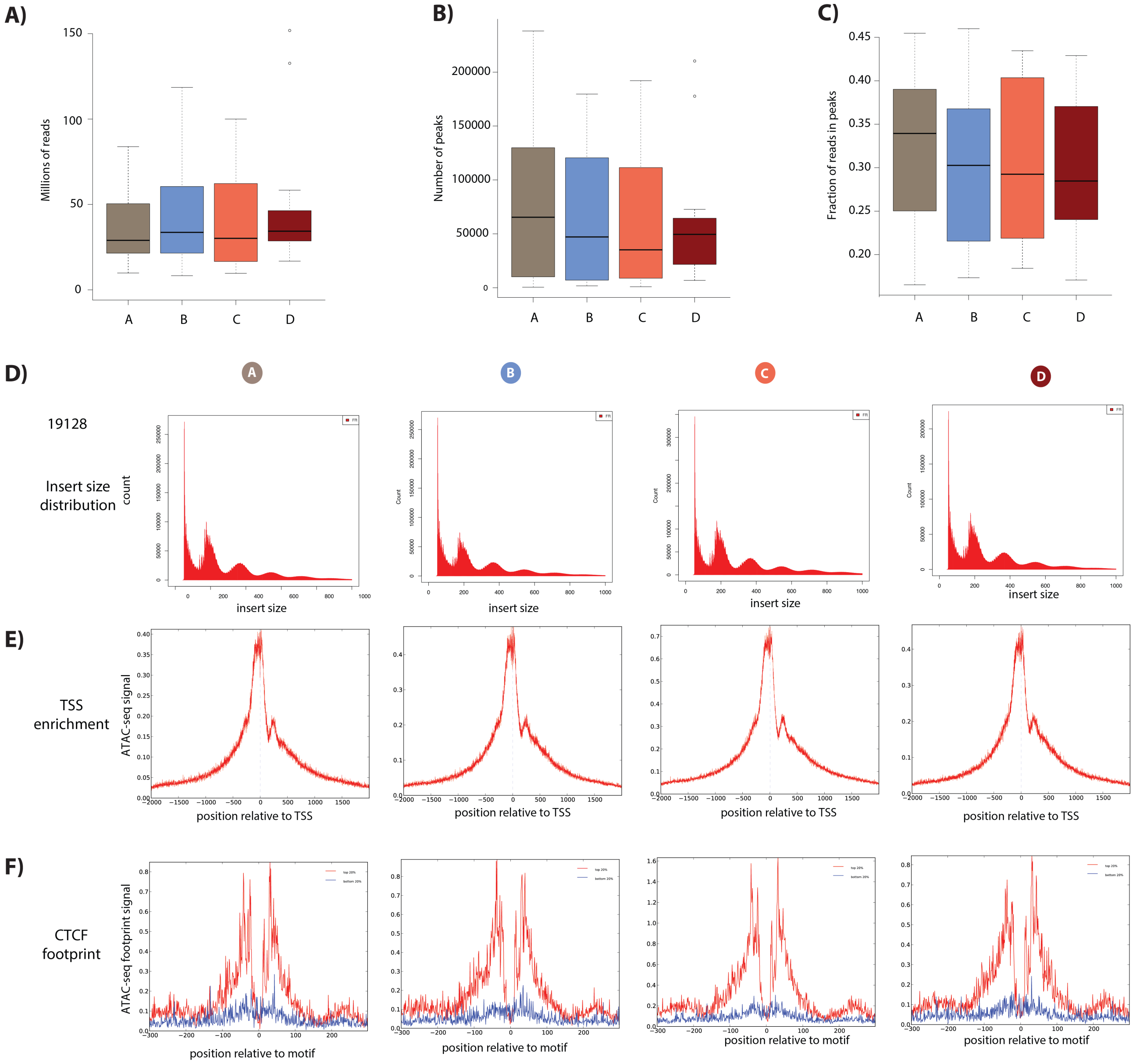

**Fig.S13: ATAC-seq library quality control. A)** The final number of ATAC-seq reads that have passed all mapping and filtering steps segregated by condition. **B)** The number of open chromatin regions (peaks) in each sample segregated by condition. **C)** The fraction of reads in peaks in all samples segregated by condition. **D)** The distribution of fragment sizes of ATAC-seq libraries from the four conditions from a representative individual (19128). **E)** The enrichment of sequencing reads around the TSS. The ATAC-seq footprint around CTCF motifs. Motifs are segregated by motif strength (top and bottom 20% of sites).

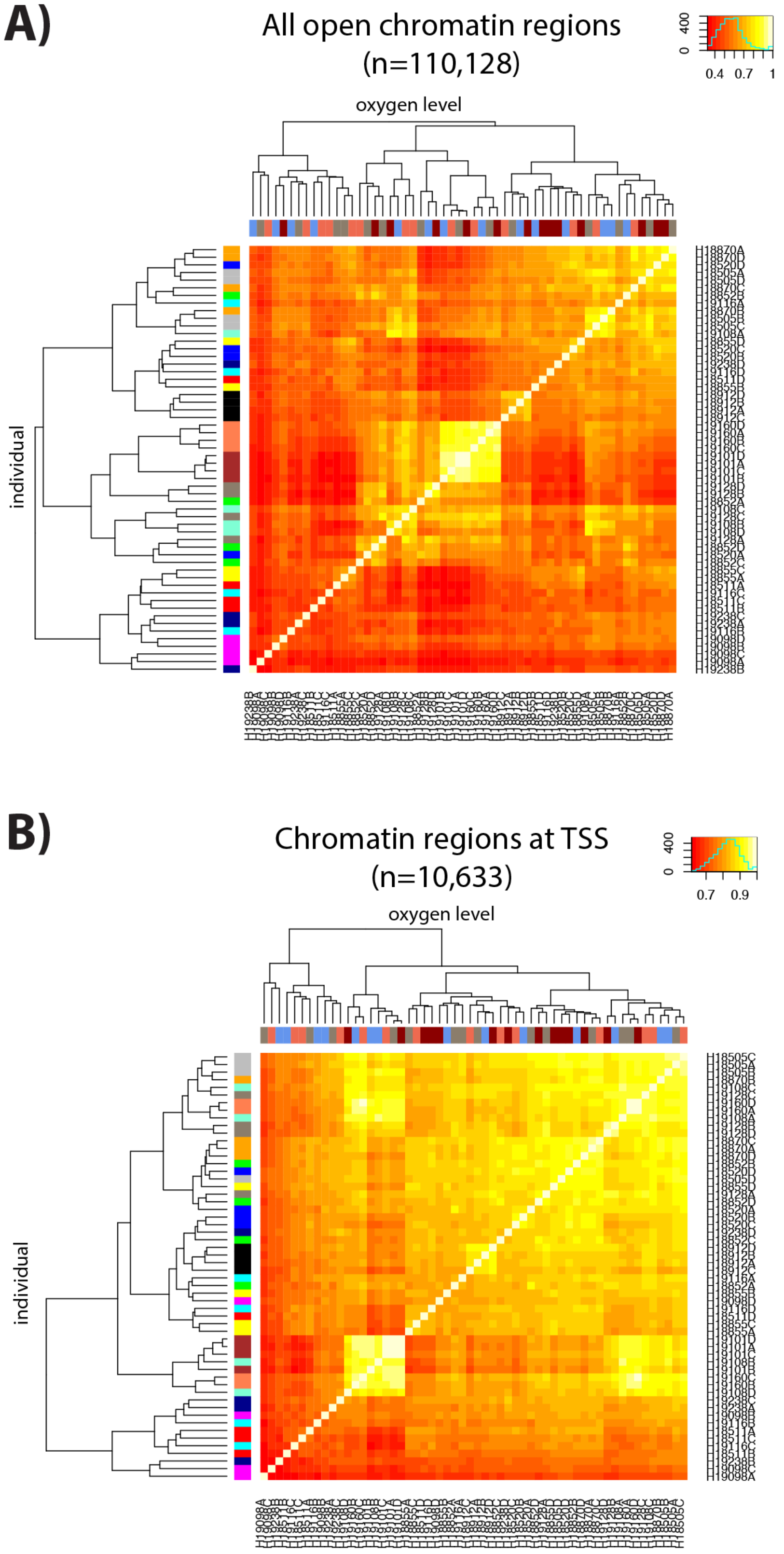

**Fig.S14: ATAC-seq libraries cluster by individual and treatment. A)** Spearman correlation of ATAC-seq read counts between samples when considering all chromatin regions. **B)** Spearman correlation of ATAC-seq read counts between samples when restricting to chromatin regions at the TSS. Colors on the x-axis refer to each of the four conditions (A: brown, B: blue, C: coral, D: red) and colors on the y-axis refer to each of the fourteen individuals.

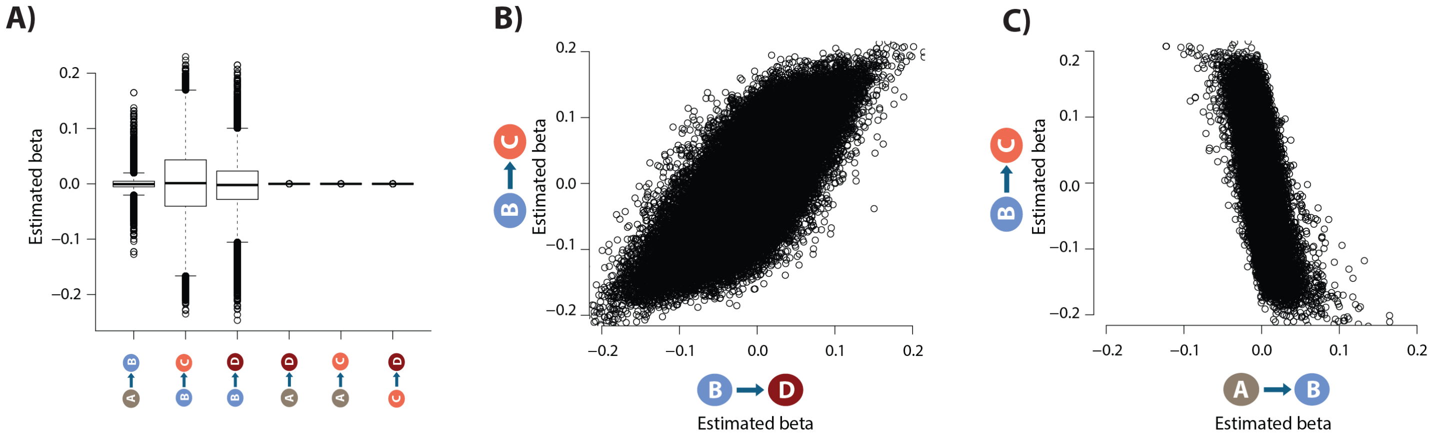

**Fig.S15: Identification of differentially accessible chromatin regions. A)** Estimated effect size (Beta) following adaptive shrinkage (ash) between each pair of conditions. **B)** Estimated Beta for each region when comparing B vs. D and B vs. C. **C)** Estimated Beta for each region when comparing A vs. B, and B vs. C.

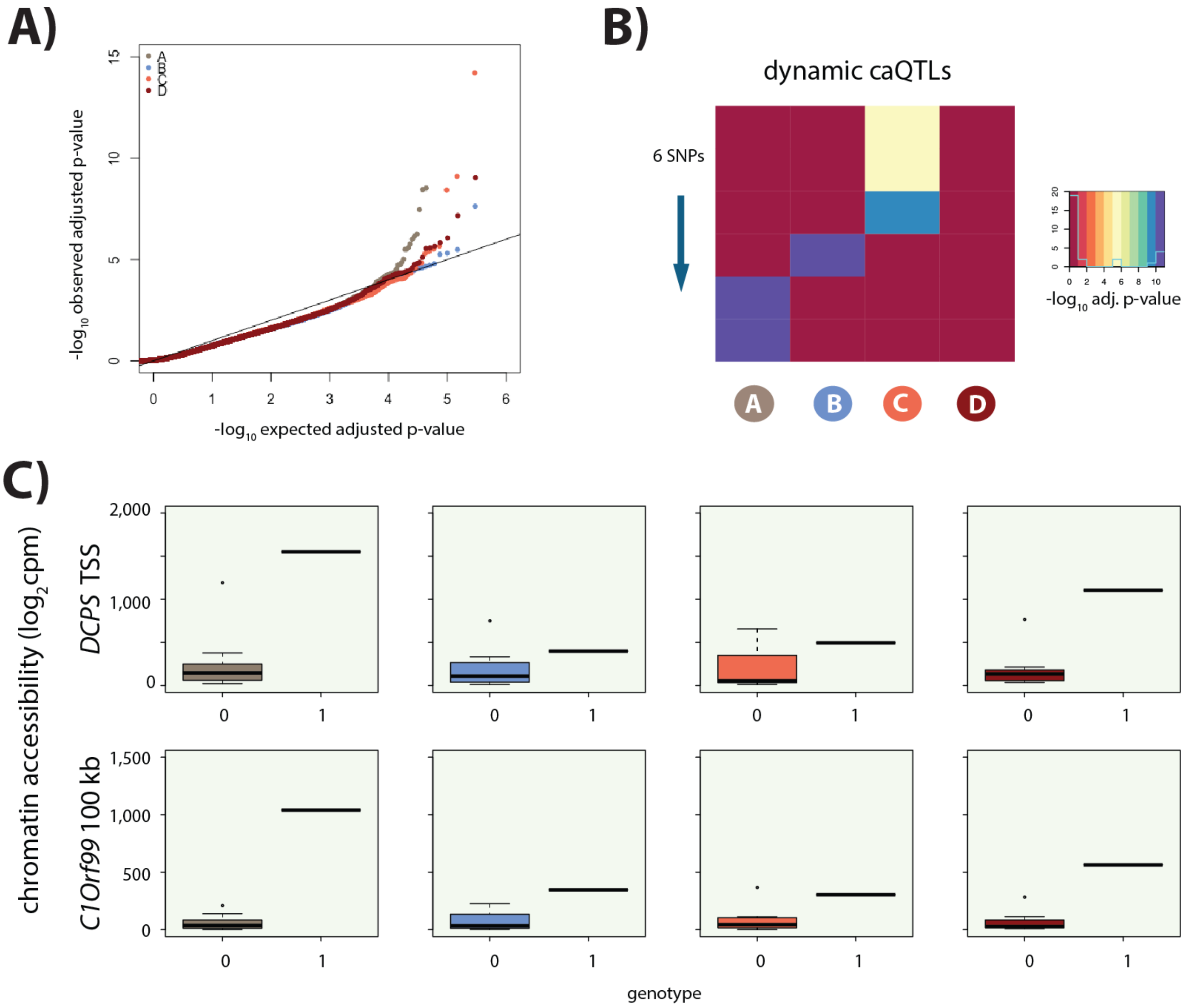

**Fig.S16: Identification of caQTLs. A)** QQ plot showing an enrichment of low p-values for the association between genotype and level of chromatin accessibility in each of the four conditions. **B)** Heatmap representing the six dynamic caQTLs. Each row represents a SNP, and the color represents the significance of the association between genotype and chromatin accessibility. **C)** Examples of two dynamic caQTLs including a region within the promoter of the *DCPS* gene, and a region ~100 kb away from the *C1Orf99* gene. The two genotype classes represented at these loci (homozygous (0) and heterozygous (1)) are included.

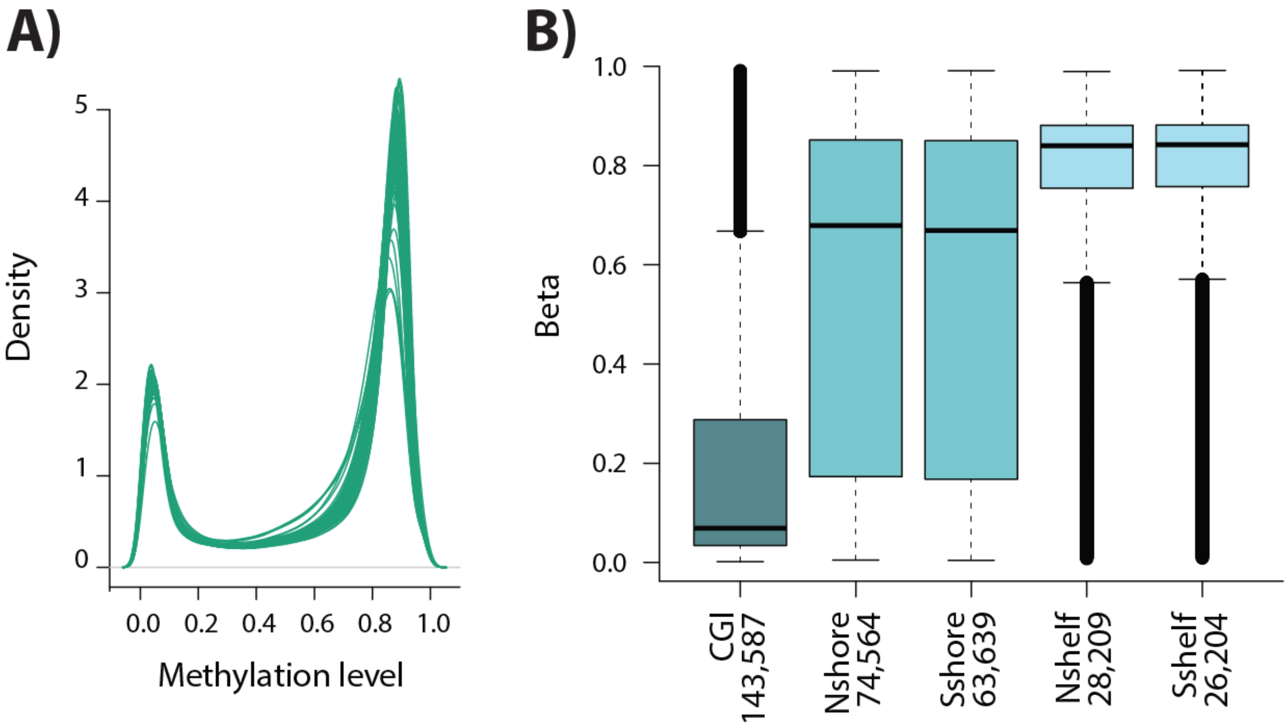

**Fig.S17: DNA methylation array quality control. A)** The distribution of Beta values across samples. **B)** The Beta level for each CpG within a CpG island (CGI), CGI north and south shores, and CGI north and south shelves.

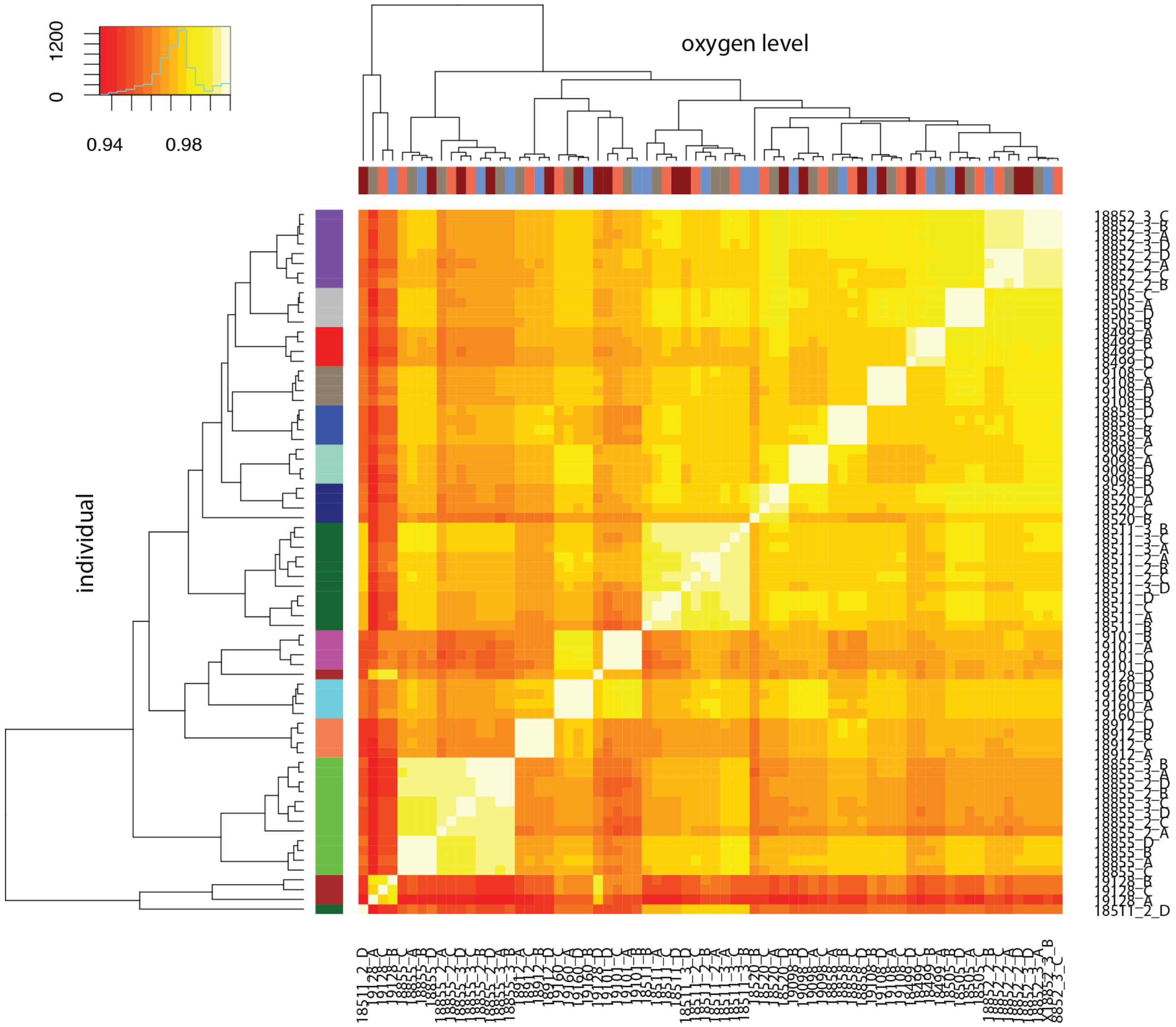

**Fig.S18: DNA methylation levels cluster by individual.** Spearman correlation of quantile-normalized Beta-values for 766,858 CpGs between all samples. X-axis represents oxygen level, and y-axis represents individual.

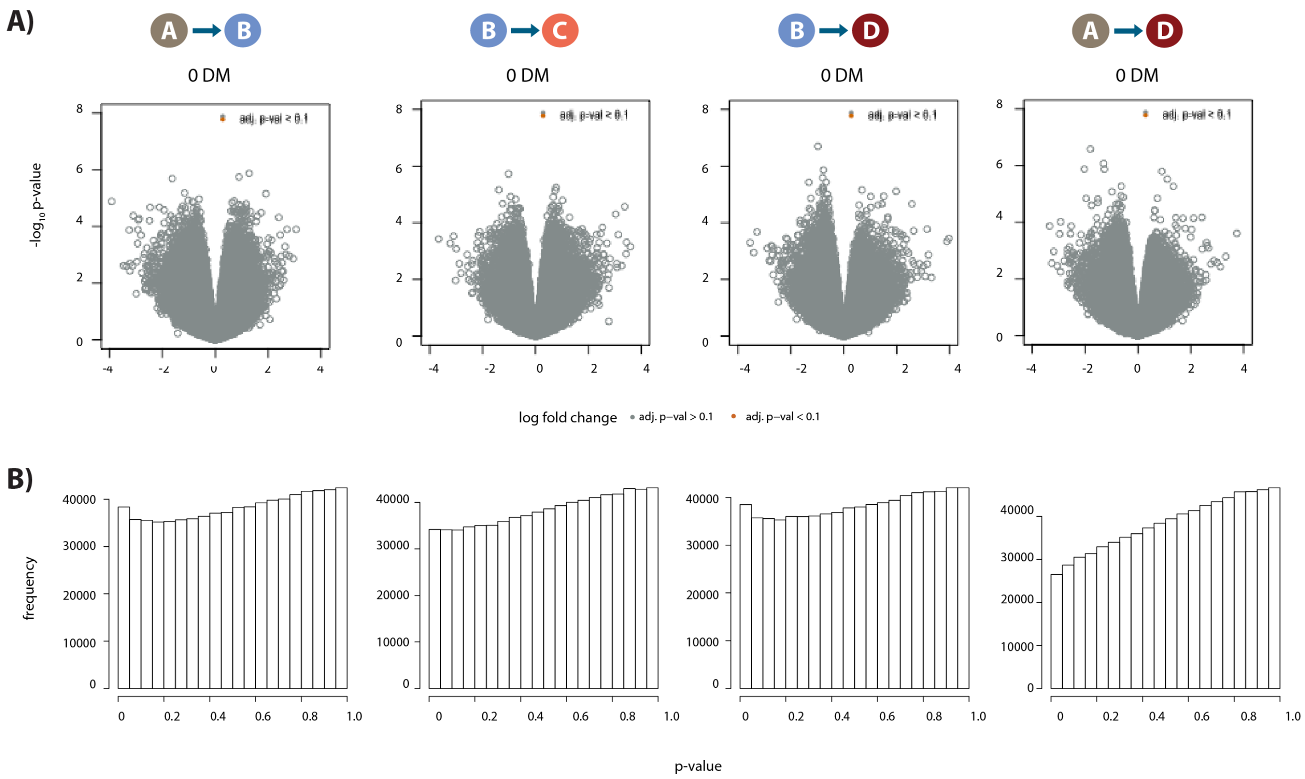

**Fig.S19: DNA methylation levels are stable across conditions. A)** Volcano plots representing the differential DNA methylation analysis across 766,658 CpGs. There are no differentially methylated (DM) CpGs. **B)** Distribution of p-values across each pair of contrasts.

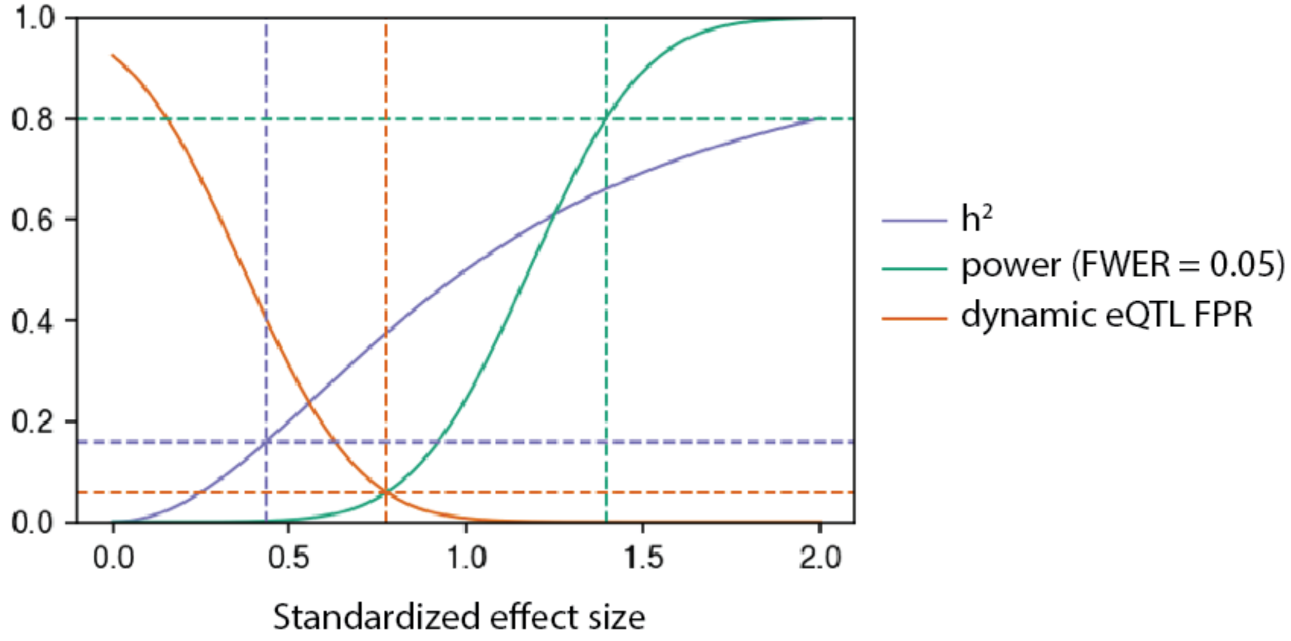

**Fig.S20: Power to detect eQTLs and false positive rate to call dynamic eQTLs.** As a function of standardized effect size: the power of this study to detect an eQTL in one condition after Bonferroni correction at level 0.05 (green), the phenotypic variance explained assuming a single causal variant ($h^2$; purple), and the false positive rate (FPR; orange) to call a dynamic eQTL assuming the true effect size is equal in all four conditions. The green dotted lines denote the effect size this study has 80% power to detect in one condition. The purple dotted lines denote the effect size corresponding to the median cis-heritability of gene expression ($h^2 = 0.16$; Gusev *et al*., 2016; Wheeler *et al*., 2016). The orange dotted lines denote the effect size for which the power to detect a QTL in one condition is equal to the FPR to call a dynamic eQTL.

**Table S1: Numbers of mapped RNA-seq and ATAC-seq reads.** The total number of reads, the number of mapped reads, the number of unbiased reads (pass allelic mapping filter), and the percentage of mapped reads for the RNA-seq data. The same metrics are shown for the ATAC-seq data as well as the total number of nuclear and unique reads, and the number of open chromatin regions (Peaks).

| | RNA-seq | | | | | ATAC-seq | | | | | |
|---|---|---|---|---|---|---|---|---|---|---|---|
| **Sample_ID** | **Total** | **Mapped** | **Unbiased** | **% mapped** | | **Total** | **Mapped** | **Nuclear** | **Unique** | **Unbiased** | **Peaks** |
| 18499_A | 13274074 | 12774871 | 8602879 | 0.96239263 | | 75898992 | 66046406 | 22888832 | 16101573 | 11433578 | 8007 |
| 18499_B | 29137307 | 27880513 | 19377082 | 0.9568665 | | 136363906 | 118388969 | 36342851 | 21732531 | 8268710 | 1974 |
| 18499_C | 17618840 | 16848276 | 11167357 | 0.95626477 | | 147109212 | 129666632 | 35889875 | 24479220 | 16624406 | 8834 |
| 18499_D | 20178367 | 19358244 | 12449962 | 0.95935632 | | 245393312 | 216720631 | 71190842 | 48428134 | 32026318 | 20592 |
| 18505_A | 61000152 | 59105803 | 44218602 | 0.96894518 | | 147548892 | 123793158 | 67785704 | 53233014 | 50404000 | 99288 |
| 18505_B | 66383475 | 64082113 | 48264602 | 0.96533231 | | 552220460 | 477651318 | 196057836 | 128921905 | 79894590 | 120504 |
| 18505_C | 73654949 | 70187145 | 52770858 | 0.95291825 | | 294509632 | 252410853 | 106254528 | 76750396 | 62240876 | 112183 |
| 18505_D | 93503767 | 89760170 | 67722802 | 0.95996314 | | 508933926 | 466266546 | 132176694 | 79952234 | 34272156 | 29367 |
| 18511_A | 87275048 | 83839277 | 61212611 | 0.96063284 | | 223280222 | 199771546 | 107499462 | 65856152 | 25487036 | 6360 |
| 18511_B | 43282709 | 41204796 | 25925729 | 0.95199208 | | 362622642 | 323669714 | 134734888 | 77526342 | 21339124 | 1780 |
| 18511_C | 65396599 | 62807010 | 43152384 | 0.96040178 | | 104707528 | 93995108 | 36821817 | 24511014 | 14486524 | 3071 |
| 18511_D | 65231093 | 62576728 | 43891079 | 0.95930829 | | 193842410 | 181414145 | 72784643 | 49555092 | 33381576 | 10345 |
| 18520_A | 155324174 | 149627180 | 110917729 | 0.96332191 | | 171193232 | 153816726 | 48869599 | 33996532 | 24817126 | 55434 |
| 18520_B | 92723262 | 89754491 | 66573746 | 0.96798246 | | 113909996 | 103490941 | 49250209 | 33541354 | 21539874 | 26223 |
| 18520_C | 80731741 | 77972317 | 55944930 | 0.96581984 | | 225670754 | 201416822 | 67937723 | 46931214 | 31939458 | 30380 |
| 18520_D | 70975926 | 68790727 | 48540194 | 0.96921211 | | 503150881 | 503150881 | 199592674 | 158987163 | 58345488 | 59592 |
| 18852_A | 20123197 | 12548026 | 6183125 | 0.62356026 | | 91975266 | 80910391 | 34626399 | 29137605 | 32509084 | 129855 |
| 18852_B | 86377262 | 80369780 | 61348638 | 0.93045065 | | 139610692 | 123969581 | 56427726 | 44608347 | 43152144 | 68011 |
| 18852_C | 83094221 | 78770813 | 57953486 | 0.94796981 | | 105440428 | 96136631 | 33210577 | 24929442 | 21804192 | 39890 |
| 18852_D | 69105112 | 65795890 | 49177751 | 0.95211321 | | 82061042 | 73583180 | 30032642 | 24939177 | 26720788 | 60950 |
| 18855_A | 44622075 | 31056755 | 22330134 | 0.69599531 | | 73412590 | 66197390 | 30381345 | 23653409 | 21467270 | 10100 |
| 18855_B | 43083433 | 41542423 | 31525741 | 0.96423196 | | 93867382 | 83857887 | 36670365 | 27073962 | 22104762 | 9827 |
| 18855_C | 95326640 | 92247773 | 69086768 | 0.96770192 | | 92329134 | 82898066 | 44621721 | 32405842 | 25088064 | 15211 |
| 18855_D | 74791880 | 72308403 | 53399182 | 0.96679483 | | 158565276 | 139042826 | 69745620 | 52688920 | 46229420 | 43799 |
| 18858_A | 24479708 | 23449164 | 15308633 | 0.95790211 | | - | - | - | - | - | - |
| 18858_B | 37816284 | 35973427 | 23690578 | 0.95126816 | | - | - | - | - | - | - |
| 18858_C | 17664628 | 16875631 | 10734974 | 0.95533464 | | - | - | - | - | - | - |
| 18858_D | 63389500 | 60619588 | 38147083 | 0.9563033 | | - | - | - | - | - | - |
| 18870_A | 83285251 | 80592918 | 57744981 | 0.96767335 | | 206701596 | 179926137 | 90702138 | 64901660 | 48947810 | 86584 |
| 18870_B | 33073289 | 31707568 | 23085282 | 0.95870622 | | 287484518 | 254534068 | 104221482 | 74674575 | 58772962 | 92168 |
| 18870_C | 43110021 | 40999695 | 30446675 | 0.9510479 | | 186428590 | 164676915 | 64609656 | 50785623 | 48740830 | 99617 |
| 18870_D | 38332664 | 36711162 | 26989210 | 0.95769921 | | 128972060 | 113397191 | 46033410 | 36228585 | 34434676 | 64467 |
| 18912_A | 84761095 | 81339751 | 57849046 | 0.95963544 | | 114728296 | 103073296 | 42774744 | 30381238 | 22269632 | 21597 |
| 18912_B | 53426335 | 50704304 | 35204067 | 0.94905076 | | 150646622 | 136928192 | 54798381 | 37895280 | 25800564 | 20046 |
| 18912_C | 75291290 | 71859868 | 49547361 | 0.95442471 | | 206380900 | 188175084 | 68272254 | 45599522 | 28383822 | 16648 |
| 18912_D | 62313003 | 59609682 | 41983277 | 0.95661706 | | 408170048 | 362928436 | 148031786 | 93497411 | 46360446 | 49318 |
| 19098_A | 64319108 | 61478974 | 42690102 | 0.95584308 | | 84805838 | 72891965 | 31229322 | 19938296 | 9877338 | 472 |
| 19098_B | 56718525 | 54252620 | 39671337 | 0.95652382 | | 326869084 | 283887723 | 120388152 | 71868062 | 25355168 | 5269 |
| 19098_C | 99513784 | 95420694 | 69762979 | 0.95886912 | | 61518746 | 53504905 | 19910386 | 13964703 | 9697158 | 935 |
| 19098_D | 94291432 | 90447815 | 65572732 | 0.95923684 | | 111045638 | 97896883 | 39106929 | 30590408 | 28617310 | 6799 |
| 19101_A | 41791627 | 40608345 | 31892100 | 0.97168615 | | 408447816 | 360350621 | 250678073 | 199132159 | 199403996 | 238073 |
| 19101_B | 44424756 | 42590301 | 33493298 | 0.95870647 | | 96278282 | 85299075 | 56468688 | 50372979 | 60498200 | 144880 |
| 19101_C | 45585076 | 44053825 | 33883532 | 0.96640894 | | 99318838 | 89512148 | 58621509 | 52805275 | 64754186 | 155535 |
| 19101_D | 54993186 | 52702068 | 40428356 | 0.95833815 | | 267006234 | 238246804 | 151538831 | 130803930 | 151857064 | 210263 |
| 19108_A | 75045688 | 72562036 | 54549304 | 0.9669048 | | 666565620 | 582192450 | 343721484 | 235061876 | 163387830 | 182451 |
| 19108_B | 122402987 | 118085837 | 90738154 | 0.96473003 | | 646218566 | 571247971 | 269071992 | 179642718 | 118543408 | 179608 |
| 19108_C | 108948403 | 105296284 | 80229311 | 0.96647845 | | 118435112 | 103365689 | 43844225 | 36602653 | 39945472 | 94177 |
| 19108_D | 75531440 | 72642872 | 53814760 | 0.96175675 | | 58027670 | 51270785 | 17411671 | 14862013 | 16800436 | 49684 |
| 19116_A | 67480144 | 65214592 | 49608734 | 0.96642639 | | 136384236 | 113613423 | 50335926 | 32298501 | 17372922 | 27377 |
| 19116_B | 44885556 | 43149149 | 32551635 | 0.96131479 | | 354799002 | 308491355 | 136910038 | 76301739 | 16149380 | 7036 |
| 19116_C | 30448184 | 29351638 | 21161665 | 0.96398649 | | 85508608 | 74170909 | 29010716 | 19442050 | 12010362 | 7061 |
| 19116_D | 37114303 | 34626358 | 25445848 | 0.93296533 | | 171251812 | 150620450 | 61635737 | 41986224 | 28094172 | 21627 |
| 19128_A | 49824910 | 47898042 | 36353716 | 0.96132722 | | 97422988 | 87659120 | 52487147 | 44259932 | 48409378 | 75274 |
| 19128_B | 89127873 | 85428483 | 67123767 | 0.95849346 | | 76628516 | 68981365 | 42202408 | 36403872 | 41523958 | 95196 |
| 19128_C | 77060247 | 72570634 | 55234011 | 0.94173892 | | 206327658 | 189834906 | 127493516 | 91998813 | 72835056 | 111412 |
| 19128_D | 61092155 | 57682764 | 43701147 | 0.94419265 | | 94083100 | 84636041 | 41641286 | 35545093 | 40160342 | 72655 |
| 19160_A | 85864447 | 82684443 | 61377326 | 0.96296483 | | 228914768 | 206224110 | 120007559 | 91187146 | 83839834 | 134015 |
| 19160_B | 54674041 | 52100580 | 38608246 | 0.95293084 | | 264357596 | 236014539 | 148012376 | 104537405 | 79486710 | 129312 |
| 19160_C | 55643747 | 53575474 | 38899078 | 0.96283009 | | 291009400 | 255529852 | 159719488 | 117754399 | 100020434 | 192041 |
| 19160_D | 48701571 | 46572060 | 32988410 | 0.95627429 | | 394585738 | 355457314 | 230980149 | 166241544 | 132661160 | 177642 |
| 18511_2_A | 17358278 | 16527759 | 11926301 | 0.9521543 | | - | - | - | - | - | - |
| 18511_2_B | 54503145 | 51980326 | 38675489 | 0.95371241 | | - | - | - | - | - | - |
| 18511_2_C | 29224912 | 27427616 | 19036688 | 0.93850123 | | - | - | - | - | - | - |
| 18511_2_D | 20747301 | 19650165 | 13668195 | 0.9471191 | | - | - | - | - | - | - |
| 18852_2_A | 43731262 | 42083993 | 29631703 | 0.962332 | | - | - | - | - | - | - |
| 18852_2_B | 17938918 | 17275639 | 12563891 | 0.9630257 | | - | - | - | - | - | - |
| 18852_2_C | 30806500 | 29648177 | 20149431 | 0.96240005 | | - | - | - | - | - | - |
| 18852_2_D | 12306206 | 11752473 | 7845269 | 0.95500376 | | - | - | - | - | - | - |
| 18855_2_A | 21248351 | 20348241 | 15001530 | 0.9576386 | | - | - | - | - | - | - |
| 18855_2_B | 33882243 | 32320306 | 24049996 | 0.95390102 | | - | - | - | - | - | - |
| 18855_2_C | 21032559 | 20084276 | 14277707 | 0.95491357 | | - | - | - | - | - | - |
| 18855_2_D | 25326814 | 24131851 | 16900384 | 0.95281827 | | - | - | - | - | - | - |
| 18511_3_A | 24403396 | 22847303 | 15578924 | 0.93623457 | | - | - | - | - | - | - |
| 18511_3_B | 33301567 | 31412705 | 23041986 | 0.94328009 | | - | - | - | - | - | - |
| 18511_3_C | 26158072 | 24437979 | 16608756 | 0.93424236 | | - | - | - | - | - | - |
| 18511_3_D | 29693799 | 28179137 | 19337816 | 0.94899063 | | - | - | - | - | - | - |
| 18852_3_A | 26010126 | 24905636 | 17305313 | 0.95753615 | | - | - | - | - | - | - |
| 18852_3_B | 48959434 | 47010551 | 34547193 | 0.96019392 | | - | - | - | - | - | - |
| 18852_3_C | 40283715 | 38595394 | 26196722 | 0.95808924 | | - | - | - | - | - | - |
| 18852_3_D | 40145008 | 38337475 | 25639178 | 0.9549749 | | - | - | - | - | - | - |
| 18855_3_A | 56943100 | 54757495 | 40867727 | 0.96161774 | | - | - | - | - | - | - |
| 18855_3_B | 39235875 | 37549457 | 28823981 | 0.95701847 | | - | - | - | - | - | - |
| 18855_3_C | 30005424 | 28691407 | 20801267 | 0.95620735 | | - | - | - | - | - | - |
